# Exploiting color space geometry for visual stimulus design across animals

**DOI:** 10.1101/2022.01.17.476640

**Authors:** Matthias P. Christenson, S. Navid Mousavi, Sarah L. Heath, Rudy Behnia

## Abstract

Color vision represents a vital aspect of perception that ultimately enables a wide variety of species to thrive in the natural world. However, unified methods for constructing chromatic visual stimuli in a laboratory setting are lacking. Here, we present stimulus design methods and an accompanying programming package to efficiently probe the color space of any species in which the photoreceptor spectral sensitivities are known. Our hardware-agnostic approach incorporates photoreceptor models within the framework of the principle of univariance. This enables experimenters to identify the most effective way to combine multiple light sources to create desired distributions of light, and thus easily construct relevant stimuli for mapping the color space of an organism. We include methodology to handle uncertainty of photoreceptor spectral sensitivity as well as to optimally reconstruct hyperspectral images given recent hardware advances. Our methods support broad applications in color vision science and provide a framework for uniform stimulus designs across experimental systems.

## 1 Introduction

From insects to primates, color vision represents a vital aspect of perception that ultimately enables a wide variety of species to thrive in the natural world. Each animal is equipped with an array of photoreceptors expressing various opsin types and optical filters that define the range of wavelengths the animal is sensitive to. But opsin expression alone does not predict how an animal might “see” colors. The downstream neural circuits mechanisms that process photoreceptor signals are critical in shaping color perception. Interrogating these neural mechanisms in a laboratory setting necessitates experimentalists to construct and present chromatic visual stimuli that are relevant to the animal in question. Outside of trichromatic primates, for which studies in human perception and psychophysics have lead the way, there is a lack of unifying methodology to assay color vision across species using disparate laboratory visual stimulation systems. Here we describe standardized methods to create chromatic stimuli, using a minimal set of light sources, that can continuously span a wavelength spectrum and be flexibly applied to photoreceptor systems in various species.

Light, the input to a photoreceptor, comprises two components: wavelength and intensity. Importantly, a photon of light of any wavelength elicits the same response once absorbed by a photoreceptor. This principle of univariance limits a single photoreceptor from distinguishing between wavelength and intensity, as different wavelength-intensity combinations can elicit the same response, rendering single photoreceptors “color-blind” (Rushton, 1972; Stockman and Brainard, 2010). By combining outputs from different types of photoreceptors in downstream neural circuits, animals can separate wavelength and intensity information to ultimately allow for color discrimination. As a result of univariance, particular wavelength-intensity combinations remain indistinguishable if they produce an equivalent set of photoreceptor responses in an animal. To the human eye, a red-green mixture is perceived as identical to pure yellow, as both of these sources equally activate the three cones. This metamerism is taken advantage of in Red-Green-Blue (RGB) screens which can display many colors using only three light sources. However, out-of-the-box RGB screens cannot easily be used to investigate color processing across animals. This is because the software that operate them are based on experimentally measured color matching functions (i.e. “matching” ratios of R/B/G to perceptual colors) that are specific to the set opsins expressed in human cones and the neural processing of their signals in the human brain. Even though such color matching functions are not available for most animals, it is still possible to leverage fundamental concepts of metamerism to construct chromatic stimuli (Fleishman et al., 1998; Tedore and Johnsen, 2017). The use of such methods has been limited, in part because of a lack of a practical framework to apply a wide range of well-established color theory concepts.

Here, we present a set of algorithms, and accompanying Python software package *drEye,* for designing chromatic stimuli which allows for the simulation of arbitrary spectra using only a minimal set of light sources. Our framework is founded on established color theory (Hempel de Ibarra et al., 2014; Kelber and Osorio, 2010; Rushton, 1972) and is applicable to any animal for which the photoreceptor spectral sensitivities are known. Our approach allows for the reconstruction of a variety of stimuli, including some that are hard to reproduce in a laboratory setting, such as natural images. Importantly, our methods are flexible to various modeling parameters and can account for uncertainties in the spectral sensitivities. Even though we provide mathematical tools to select appropriate light sources, our methods are ultimately agnostic of the hardware used for visual stimulation. For this reason, our method can be used as a color management tool to control conversion between color representations of various stimulus devices. We illustrate basic principles as well as examples of our algorithms using the color systems of mice, bees, humans, fruit flies, and zebrafish.

## 2 Color theory and color spaces

As color scientists, we aim to understand how a given animal processes spectral information and thus perceives color. A “perceptual color” space gives an approximation of how physical properties of light are experienced by the viewer.

CIE 1931 color spaces, for instance, are defined mathematical relationships between spectral distributions of light and physiologically perceived colors in human vision (Smith and Guild, 1931). These spaces were derived from psychophysical color matching experiments (Smith and Guild, 1931; Wright, 1929), and are an essential tool when dealing with color displays, printers and image recording devices. Because the underlying quantitative transformation from the spectral distributions of light to the perceived colors in humans is dependent on both the cone spectral sensitivities and the neural mechanisms that process them, human color spaces do not transfer to other animals.

Can we approximate the perceptual space of animals using available physiological and/or behavioral information? A perceptual color space is the result of a series of transformations starting from the stimulus itself. The stimulus can be represented in “spectral space”, simply describing the spectral distribution of light. This space however is high dimensional, and therefore difficult to work with. Instead, a lower dimensional color space can be constructed, by taking into account the photoreceptor spectral sensitivities of the viewer. A photoreceptor’s spectral sensitivity defines the relative likelihood of photon absorption across wavelengths (e.g. Fig. 1A-C and S1A-B). Weighting a total light power spectrum by the photoreceptor’s spectral sensitivities renders *n* effective values of a stimulus – with *n* being the number of photoreceptor types. The n values compose a *n*-stimulus specification of the objective color of the light spectrum for an animal, called the photon capture. This results in a *n*-dimensional receptor-based “capture space”. For dichromats such as new world monkeys or mice, this receptor-based space is two-dimensional (Fig. 1D). For trichromats, such as humans or bees, it is three-dimensional (Fig. 1E) and for tetrachromats, such as zebrafish and fruit flies, it is four-dimensional (Fig. 1F).

**Figure 1.**
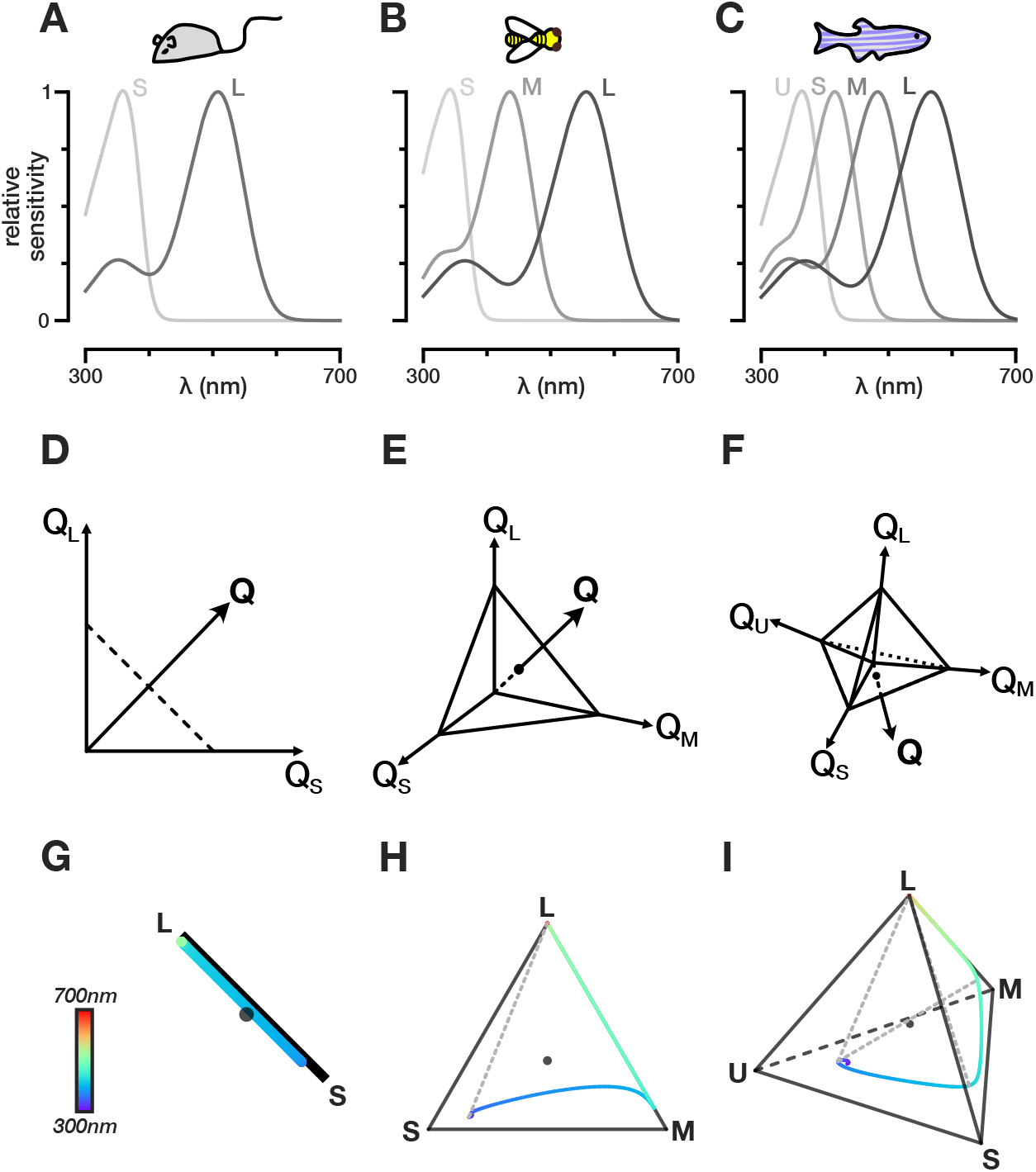
Color and chromatic spaces of di-, tri-, and, tetrachromatic animals. **(A-C)** Spectral sensitivity functions for the different opsins expressed in the photoreceptors of the mouse, the honey bee, and the zebrafish, respectively. Photoreceptors assigned the label long (L), medium (M), short (S), ultrashort (U) from the longest to shortest wavelength-sensitive photoreceptor. **(D-F)** Schematic representation of receptor-based color spaces of di-, tri-, and, tetrachromatic animals, respectively. *Q* denotes capture. **(G-I)** Chromatic diagrams for the mouse, the honey bee, and the zebrafish, respectively. The colored line indicate the loci of single wavelengths in the chromatic diagram. The dotted lines indicate hypothetical non-spectral color lines that connect the points along the single wavelength color line that maximally excite non-consecutive photoreceptors.

In addition, within this *n*-dimensional receptor-based capture space, it is often useful to define a hyperplane, where vector points sum up to 1, and where color is therefore represented independently of intensity. The resulting “chromaticity diagram” is the n – 1 simplex where a point represents the proportional capture of each photoreceptor (Fig. 1G-I and Fig. S1C-D). For dichromats this visualization simplifies to a line, for trichromats, it is a triangle and for tetrachromats, a tetrahedron. The loci of single wavelengths can be mapped onto these spaces as a one-dimensional manifold, as can theoretical “non-spectral” color lines. Non-spectral colors result from the predominant excitation of photoreceptor pairs that are not adjacent along the single wavelength manifold (Stoddard et al., 2020; Thompson et al., 1992).

Although the number of photoreceptors, which determines the dimensionality of the receptor-based space, is not always equal to the effective dimensionality of perceived colors (Jacobs, 2018), it provides its theoretical maximum. The effective dimensionality depends on the processing of photoreceptor signals in the brain. In fact neural processing effectively distorts the shape of receptor based spaces, to eventually produce a perceptual space, where distances do not necessarily match the distances measured in receptor-based spaces.

Receptor-based spaces are however a good starting point to mathematically approximate the transformations that the brain applies to photoreceptor inputs. They can in particular serve to design relevant chromatic stimuli to interrogate these transformations experimentally. Throughout this paper, we will use receptor-based color spaces as the foundation for a unified framework for developing such chromatic stimuli.

## 3 Reconstructing arbitrary light spectra: A general framework

Probing an animal’s color vision requires measuring behavioral or physiological responses to relevant chromatic stimuli. Amongst these are artificial stimuli which are constructed to probe specific aspects of visual processing, such as a set of Gaussian spectral distributions to measure spectral tuning, naturalistic stimuli, such as measured natural reflectances, or randomly drawn spectral stimuli, akin to achromatic noise stimuli. Current methods to display such stimuli often do not take into account the visual system of the animal under examination, and instead focus on spectral space which is often not as relevant functionally. Instead, we have developed a method that allows for the reconstruction of a wide range of chromatic stimuli, with only a limited number of light sources, that can be applied across animals for which spectral sensitivities are known. Here we describe the core method for light spectra reconstruction, followed by highlights of important considerations regarding the stimulus system and aspects of the fitting procedure. An overview of the method is illustrated in Figure 2 for an idealized dichromatic animal chosen for ease of visualization.

**Figure 2.**
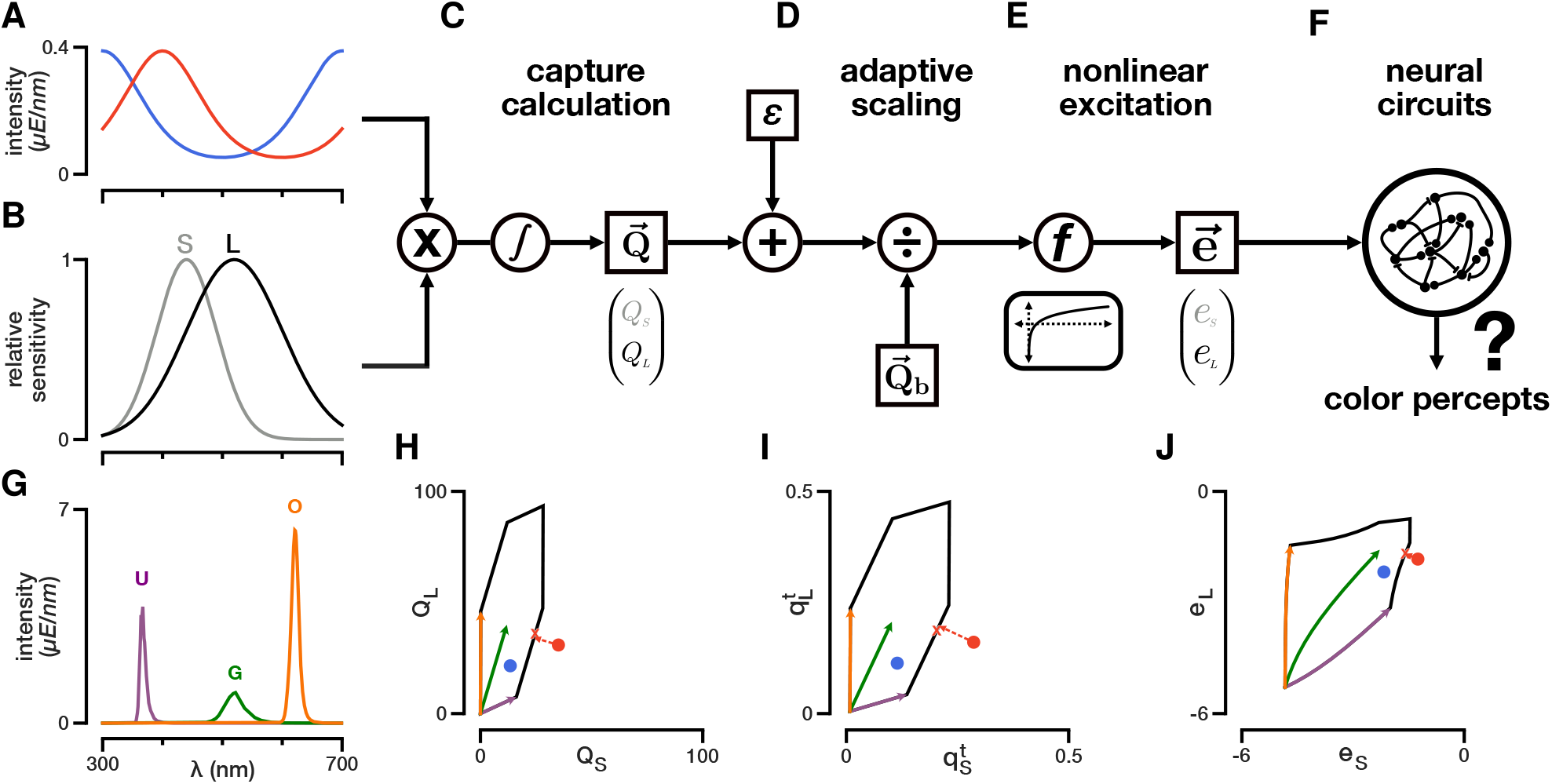
Schematic representation of the photoreceptor model. **(A)** Two example spectral distributions of light constructed artificially. Red: exp {sin(2*π*(λ — 300*nm*)/400*nm*)}; blue: exp {cos(2*π*(λ — 300*nm*)/400*nm*)}. **(B)** Artificial spectral sensitivities constructed using a Gaussian distribution with mean 440nm and 520nm and standard deviation 50nm and 80nm for the shorter (S) and longer (L) wavelength-sensitive photoreceptor, respectively. **(C)** To calculate capture, the lights in A hitting the photoreceptors in B are each multiplied by the spectral sensitivities of each photoreceptor and integrated across wavelengths. A small baseline capture value *ϵ* can be added to the light-induced capture value. **(D)** To calculate the relative capture, the absolute capture calculated in C is divided by the background capture according to *von Kries* adaptation. **(E)** A nonlinear transformation is applied to the relative capture values to obtain photoreceptor excitations. **(F)** Photoreceptor signals are further processed in downstream circuits to give rise to color percepts. **(G)** Example stimulation system consisting of a set of three LED light sources at their maximum intensity (violet, green, and orange). **(H-J)** Capture space, relative capture space, and excitation space of photoreceptors in B. The colored vectors represent the integration of the LED spectra in G with the spectral sensitivities in B. The colors match the colors of the LEDs in G. These vectors can be combined arbitrarily up to their maximum LED intensities and define the gamut of the stimulation system (black lines). The red and blue circles are the calculated captures, relative captures, and excitation values for the spectra in A, respectively. The red-colored spectrum is out-of-gamut for the stimulation system defined in G. Projection of this out-of-gamut spectrum onto the gamut of the stimulation system gives different solutions when done in capture, relative capture, or excitation space (red line). The red X drawn at the edge of the stimulation system’s gamut corresponds to the projection of the red-colored spectrum onto the gamut in excitation space (i.e. the fit in excitation space).

### 3.1 Building receptor-based color spaces

The light-induced photon capture *Q* elicited by any arbitrary stimulus *j* is calculated by integrating its spectral distribution *I_j_* (λ) (in units of photon flux: *E* = *mol/s/m*^2^) with the effective spectral sensitivity *S_i_* (λ) of photoreceptor *i* across wavelengths (Fig. 2A-C). Even when no photons hit a photoreceptor, it randomly produces dark events (Barlow et al., 1993; Chu et al., 2013; Stockman and Brainard, 2010). Mathematically, we can add these dark events as a baseline capture *ϵ_i_* to the light induced capture. By multiplying this sum by the absolute sensitivity of photoreceptor *i* (*C_i_*), we obtain the total absolute capture 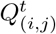:

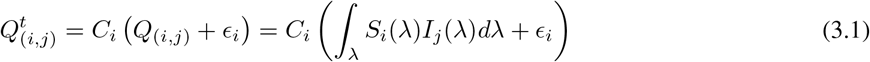

When we calculate the total capture for all *n* photoreceptor types present in an animal, we get a vector that can be represented as a point in the receptor-based capture space (Fig. 2C and H):

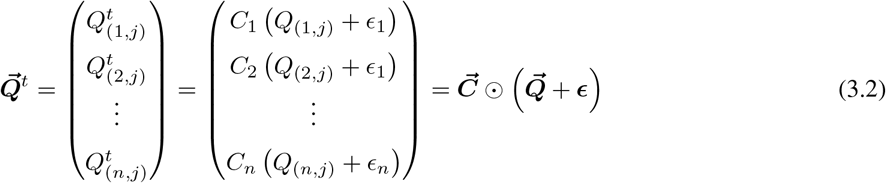

Equations 3.1 and 3.2 assume we know the spectral sensitivity of each photoreceptor and two more quantities: the absolute sensitivity *C_i_* and the baseline capture *ϵ_i_*. Unlike the spectral sensitivities of photoreceptors, both *C_i_* and *ϵ_i_* are usually unknown (and difficult to estimate) for most model organisms. In many conditions, it is assumed that the photoreceptors adapt to a constant background light according to *von Kries* adaptation (Kelber et al., 2003; Stockman and Brainard, 2010). This removes *C_i_* from the equation, and we obtain the relative light-induced capture *q*_(*i,j*)_ and baseline capture *η*_(*i,b*)_ for background *b*:

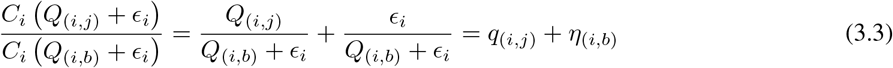

For all *n* photoreceptor types, we obtain a vector (**q**^*t*^ = **q** + *η*) representing a point in relative receptor-based capture space (Fig. 2D and I). Note that equation 3.3 is mathematically equivalent to setting *C_i_* to 1/ (*Q*_(*i,b*)_ + *ϵ_i_*). Thus, the relative photon capture is simply a form of multiplicative scaling that has been shown to approximate adaptational mechanisms within isolated photoreceptors (Clark et al., 2013; Juusola, 1993; Stockman and Brainard, 2010).

Finally, if we assume that the light-induced capture is much larger than the baseline capture, we can drop **η** so that **q** = **q**^*t*^. However, we will show in a later example why setting a baseline capture value to a specific low value can have practical uses for designing color stimuli even when we lack knowledge of the exact biophysical quantity ascribed to it. Receptor-based photon capture spaces do not take into account the neural transformation applied by the photoreceptors themselves once photons are absorbed to give rise to electrical signals. It can therefore be beneficial to further convert our relative capture values to photoreceptor excitations **e** by applying a transformation *f* that approximates the change in the response in photoreceptors (Fig. 2E and J):

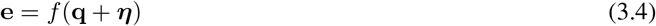

Common functions used for animal color vision models are the identity, the log, or a hyperbolic function (Chittka, 1992; Clark et al., 2017; Hempel de Ibarra et al., 2014; Vorobyev and Osorio, 1998). Applying any of these functions – except the identity function – will change the geometry of the color space and thus distances measured between points (Fig. 2H-J). If the transfer function is not known for an animal’s photoreceptors, the identity function (i.e. the linear case) can be used or a transfer function can be reasonably assumed given measured transfer functions in other animals or photoreceptor types. Photoreceptor excitation values are the de facto inputs to the visual nervous system of the organism. We will therefore consider spectral stimuli within photoreceptor excitation spaces as a foundation for our subsequent fitting procedures.

### 3.2 Fitting procedure

The goal of our method is to enable experimenters to use only a limited set of light sources to create metamers that “simulate” arbitrary light spectra for a given animal or animals. In order to do so, we use a generalized linear model of photoreceptor responses (Fig. 2A-E) to adjust the intensities of a set of the light sources in order to map intended spectral distributions onto calculated excitations of each photoreceptor type. Using equations 3.1–3.4, we can calculate photoreceptor excitation for any desired light stimulus. This results in an *n*-dimensional vector e that represents the effect of this visual stimulus on the assortment of photoreceptors of the animal. Instead of presenting this particular arbitrary distribution of light, we can use a visual stimulus system composed of a limited set of light sources to approximate this vector e and thus match the responses to our desired light stimulus. In figure 2, this corresponds to finding a coefficient for each light source vector to approximate the coordinates of the visual stimulus points in the 2D excitation space. This operation could theoretically be done in capture or relative capture space. However given that each transformation distorts the distances between points, it is more appropriate to perform this operation in excitation space, given that it is closer to perceptual space.

Given an animal’s *n* photoreceptors and *m* available light sources, we can construct a normalized capture matrix **A** using equation 3.3:

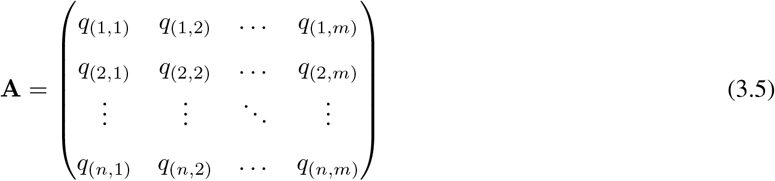

Here, *q*_(*i,j*)_ is the relative light-induced photon capture of photoreceptor *i* given the light source *j* at an intensity of one unit photon flux. Calculating **A** requires knowledge of the spectral distribution of each light source, which can be obtained using standard methods in spectrophotometry (Franke et al., 2019; Heath et al., 2020). This will also yield the intensity bounds of each light source. We denote the lower bound intensity vector as *ℓ* and the upper bound intensity vector as *u*. The theoretical minimum value for *ℓ* is 0 as a light source cannot show negative intensities.

To match the desired photoreceptor excitations, we need to find the intensity vector x for the available light sources so that the calculated excitations of the system match the desired photoreceptor excitations e:

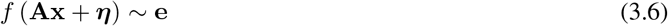

To find the optimal x, we first consider two points. First, x needs to be constraint by the lower bound *ℓ* and upper bound u. If an experimenter wants to find the best fit independent of the intensity range of the stimulation system, we only need to have a non-negativity constraint for x (theoretical minimum of *ℓ*). Second, an experimenter may want to weight each target photoreceptor excitation differently, if certain photoreceptors are thought to be less involved in color processing (see e.g. Heath et al. (2020)). Thus, we obtain a constrained objective function that minimizes the weighted (w) difference between our desired excitations (e) and possible excitations (*f* (**A**x + *η*)) subject to the intensity bound constraints *ℓ* and *u*:

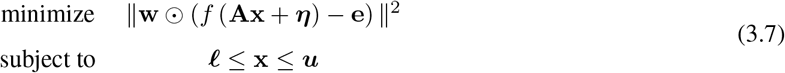

To ensure that we consistently find the same x, we use a deterministic two-step fitting procedure. First, we fit the relative capture values – before applying any nonlinear transformation – using constrained convex optimization algorithms. Next, we initialize x to the value found during linear optimization and use nonlinear least squares fitting (Trust Region Reflective algorithm) to find an optimal set of intensities that match the desired excitations. This two-step fitting procedure is deterministic and ensures that the nonlinear solution found is the closest to the linear solution, if the nonlinear optimization problem is not convex. If the transformation function is the identity function, the second step is skipped completely as the first step gives the optimal solution. There are a few important points to note regarding this fitting procedure, which we address below.

### 3.3 Gamut and visual stimulation systems

So far we have not considered the hardware and the ability of a visual stimulus system to represent colors. In human color vision, the “gamut” represents the total subset of colors that can be accurately represented by an output device, such as a LED stimulation system (Balasubramanian and Dalal, 1997). In order to generalize this concept and use it to design stimulus systems that are adequate for our fitting procedures, we have derived a gamut metric that corresponds to the “percentage of (animal) colors reproduced by a stimulation system”. To calculated this metric we separately consider a “perfect” stimulation system, where the intensity of each unit wavelength along the (animal) visible spectrum can be varied independently, and a “real” stimulation system, composed of a combination of light sources. We derive a measure of size that each system occupies in an animal’s corresponding chromaticity diagram and calculate their ratio (See methods for details). This mathematical tool can be applied to any set of light sources for which the spectra have been measured, and can be used to optimally select a set of light sources in the context of our fitting procedure.

For illustration purposes we consider a set of commercially available LEDs (Fig. S2A-B) which can be combined to create a stimulus system (Heath et al., 2020). We vary the composition and number of LEDs of a stimulus system, and calculate the metric for mice, bees, humans, fruit flies and zebrafish. We find that if the number of LEDs is below the number of photoreceptors, each LED added to the system significantly increases the fraction of colors that can be represented (Fig. S2H-L). Adding more LEDs than this only minimally improves the system (Fig. S2H-L). Examining the distribution of all *n*-sized LED stimulus systems (with n being the number of photoreceptor types of the animal) highlights that different animals allow for more or less freedom of LED choice (Fig. S2M-Q). Interestingly, LED combinations that would be chosen according to the peak of the sensitivities, a commonly used strategy when designing stimulus systems (Schnaitmann et al., 2018; Zimmermann et al., 2018), most often are not included in the 10% largest gamuts of all *n*-LED combinations (Fig. S2M-Q). This is due to the fact that our metric takes into account the shape and overlap of the sensitivities and LEDs, in addition to the peak of the sensitivities and LEDs.

Finally, a desired property of a given stimulus system may be to enable experiments across vastly different intensity regimes. As stimulus intensities are increased, LEDs will reach their maximal intensities (Fig. S2B) and the gamut of the stimulation system will decrease (Fig. S2H-L). At higher intensities of a stimulus, adding additional LEDs can enable reconstruction of more colors. This gamut metric is therefore a useful tool for assessing the suitability of an existing visual stimulation system or selecting light sources for *de novo* assembly.

### 3.4 In and out of gamut fitting

A desired capture value of light can be within the gamut of the stimulation system or out-of-gamut (e.g. Fig. 2H). If the desired captures of a stimulus set are within the gamut of the stimulus system, applying any excitation transformation or changing the weighting factor w will have no effect on the fitted intensities as an ideal solution exists (Fig. 2H-J). In this case, the second step of the fitting procedure (i.e. the nonlinear optimization) will be skipped to improve efficiency. Conversely, the intensities found when fitting captures outside the system’s gamut can vary depending on the chosen non-linearity and weighting factor w (Fig. S3). In these cases, it is especially important to consider the light conditions during experiments (photopic, mesopic, scotopic). According to various models of photoreceptor noise, the noise of photoreceptors is constant in dark-adapted conditions and becomes proportional to the capture in light-adapted conditions (Weber’s law) (Chen et al., 2012; Stockman and Brainard, 2010). Thus the monotonic transformation function *f* chosen for each condition should be the identity or the log, respectively, in order to ensure homogeneity of variance. We have found that using a log transformation and adding a small constant baseline capture ε provide a good prediction of photoreceptor responses across intensities for the fruit fly in the dark-adapted state (Fig. S4). This nonlinearity effectively rectifies the calculated captures for small values that are indistinguishable from dark and smoothly transitions between a linear and logarithmic regime. Furthermore, this transformation approximates the measured responses of other photoreceptors (Kawasaki et al., 2015) and prevents a zero division error in dark-adapted or close to dark-adapted conditions when using a log transformation.

### 3.5 Gamut correction prior to fitting

For humans, many displays use gamut correction algorithms to adjust how out-of-gamut colors are represented (Bae et al., 2010). For example, an image that is too intense will be scaled down in overall intensity in order to fit within the gamut. This can also be achieved with our method by fitting the image without any upper intensity bounds and then rescaling the fitted intensities to fit within the gamut of the stimulation system. Alternatively, capture values can be scaled prior to fitting, so that they are within the intensity bounds of the stimulation system (Suppl. C for details). For values that are completely outside of the color gamut – they cannot be reproduced by scaling the intensities – values are usually scaled and clipped in a way to minimize “burning” of the image (Bae et al., 2010). An image is burned when it contains uniform blobs of color that should have more detail. procedures to minimize burning of an image for humans usually involve preserving relative distances between values along dimensions that are most relevant for color perception. But these procedures are imperfect and will ultimately distort some of the colors. To generalize such procedures to non-primate animals, we have implemented an algorithm that assesses which capture values are outside of a system’s gamut and adjusts the capture values across the whole image to minimize “burning-like” effects by preserving relative distances between target values (Suppl. C for details). Our gamut-corrective procedures can be applied before fitting or optimized during fitting using our package. Applying gamut-corrective procedures will ensure that the relative distribution of capture values of the fitted image resembles the distribution of the original image, thus minimizing burning-like effects. Additionally, gamut correction does not require a specification of the nonlinear transformation function as all capture values are projected into the system’s gamut (as discussed in section 3.4).

### 3.6 Underdetermined stimulus system

If the stimulus system is underdetermined – i.e. there are more light sources to vary than there are types of photoreceptors – then a space of target excitations can be matched using different combinations of intensities (Fig. S3). By default, the intensities that have smallest L2-norm within the intensity constraints are chosen. This will choose a set of light source intensities that generally have a low overall intensity and similar proportions. However, in our package *drEye*, we provide alternative options such as maximizing/minimizing the intensity of particular light sources or minimizing the differences between the intensities of particular light sources. An underdetermined system can also be leveraged in other ways, as we discuss in section 5.

## 4 Example applications

### 4.1 Targeted stimuli

To illustrate our approach, we have applied our fitting procedure to two sets of targeted stimuli using the five example animals and LED-stimulation systems comprised of the LEDs introduced previously (Fig. S2A-B). The first stimulus is a set of Gaussian spectral distributions. Simulating this set of stimuli is similar to exciting the eye using a monochromator and mapping responses along a one-dimensional manifold in color space. We have simulated such a set of stimuli previously to map the tuning properties of photoreceptor axons in the fruit fly (Heath et al., 2020). The second stimulus is a set of natural reflectances of flowers multiplied with a standard daylight spectrum (Chittka, 1997; Chittka et al., 1994; Gumbert et al., 1999; Hernández-Andrés et al., 2001) (Fig. 3A). In both cases, we assume that photoreceptors are adapted to the mean of either stimulus set. We include a small baseline *ϵ* of 10^-3^ *μ*E and a log transformation in our photoreceptor model. Each photoreceptor type is equally weighted. To compare the results for different stimulation systems, we calculated their R^2^ values as a measure of goodness-of-fit. For the single wavelength stimulus set, a good fit is usually achieved with a *n*-size LED stimulus system, although for tetrachromatic animals some wavelengths may require more LEDs for a good reconstruction (Fig. 3B-D and H-J). Furthermore, more LEDs become necessary for high-intensity simulations (Fig. S5A-F). For the natural reflectance stimulus set, a *n*-size LED stimulus system usually gives a perfect fit, if an appropriate LED set is chosen (Fig. 3E-G and H-J). However, more LEDs than this improve fits for high-intensity spectra (Fig. S5A-F). The naturalistic stimulus set is more correlated across wavelengths. Thus, these types of stimuli are usually easier to simulate given different stimulation systems, as they are covered by a stimulus system’s gamut – especially when adapted to the mean.

**Figure 3.**
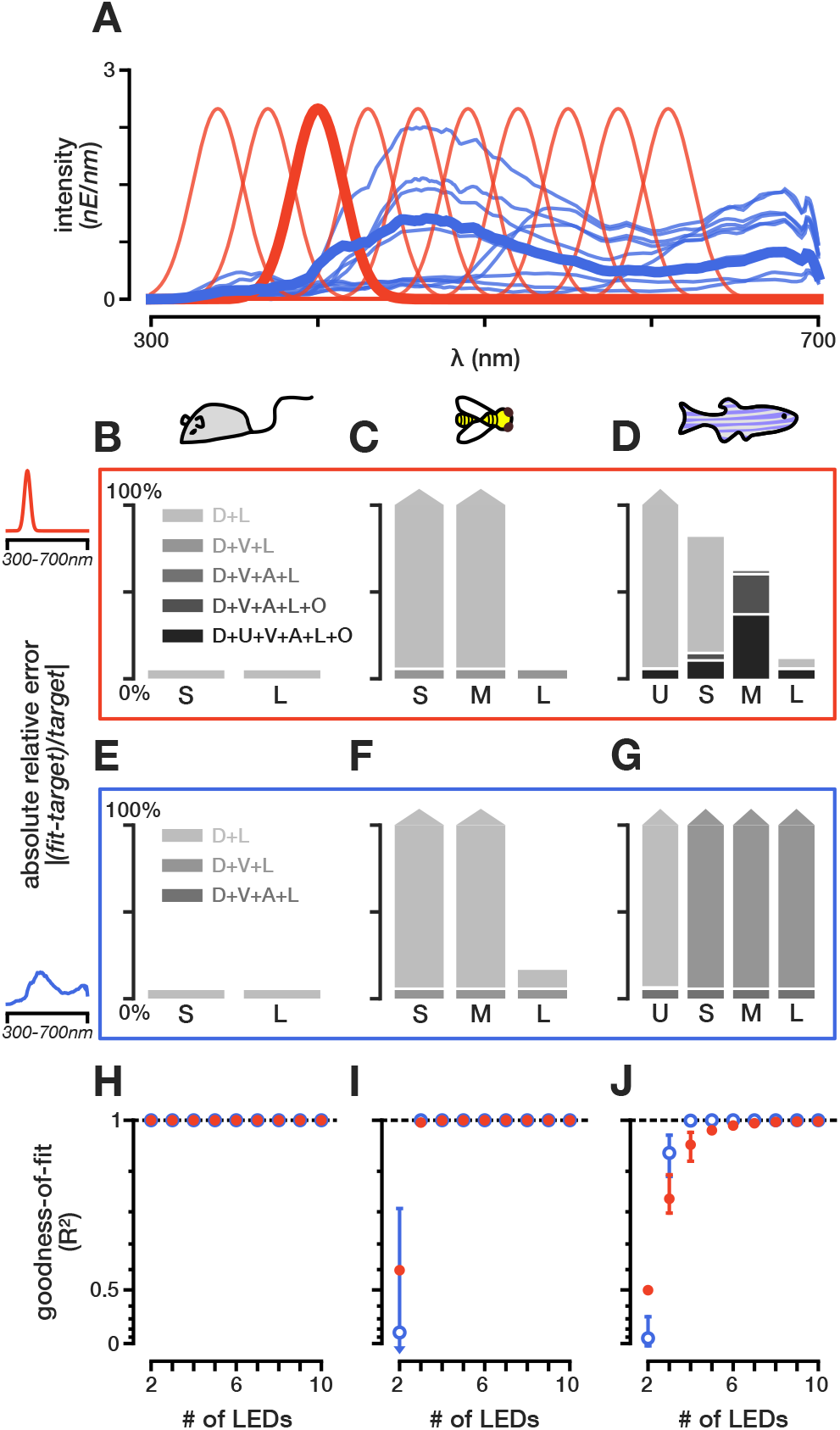
Fitting targeted stimuli to different model organisms. **(A)** Example target spectra to be reconstructed: a set of natural spectral distributions (blue) and a set of Gaussian spectral distributions (red). **(B-D)** Absolute relative error of fitting the 400nm spectrum to the mouse, honey bee, and zebrafish, respectively. For the mouse two LEDs are sufficient to recreate the spectrum, but for the zebrafish a perfect recreation is not even possible with six LEDs. **(E-G)** Absolute relative error of fitting a natural spectrum to the mouse, honey bee, and zebrafish, respectively. For the mouse two LEDs, for the honey bee three LEDs, and for the zebrafish four LEDs are sufficient to perfectly simulate the spectrum. **(H-J)** Goodness-of-fit (R^2^) values for the best LED sets (top 10%) across different number of LED combinations for the mouse, honey bee, and zebrafish, respectively. The barred lines for each point correspond to the range of *R*^2^ values achieved for the top 10% of LED combinations. The y-axis is plotted on an exponential scale to highlight differences in the goodness-of-fit close to 1.

### 4.2 Random stimuli

So far, we have assumed to have a set of spectral distributions to simulate. However this is not necessarily the case. Random sampling of a color space can be a useful way to probe a chromatic system. Similar to stimulating the eye using artificial achromatic stimuli, such as random square or white noise stimuli (Meyer et al., 2017; Zhuang et al., 2017), we can stimulate the eye using artificial chromatic stimuli to extract detailed chromatic receptive fields. This can theoretically be done in either spectral- or receptor-based color space. To compare both methods, we will consider the excitation space of the medium- and long-wavelength photoreceptors of our five animals – the mouse, honey bee, zebrafish, human, and fruit fly. For this example, the stimulus system consists of the violet and lime LEDs (Fig. S2A). Photoreceptors are adapted to the sum of 1*μE* photon flux of light for both LEDs, and the photoreceptor model incorporates a small *ϵ* of 10^-3^*μE* and a log transformation. We sample 121 individual stimuli that equally span a two-dimensional plane from −1 to 1. The samples are drawn either from relative LED intensity space – log(**i**/**i**_*b*_) with **i** being the LED intensities and **i**_*b*_ being the background intensities – or from photoreceptor excitation space. In the latter case, we fit LED intensities using our fitting procedure to best match the desired excitations. Points outside of the system’s gamut will be clipped as per the fitting procedure. No fitting is required when directly drawing LED intensities. When drawing equally spaced samples in relative LED space, the samples are highly correlated along the achromatic dimension of the excitation space (i.e. *e_L_ = e_M_*) and do not span the available gamut in excitation space (Fig. S6A-C). Consequently, any neural or behavioral tuning extracted from such a stimulus set can be biased towards specific directions in color space (Meyer et al., 2017; Weller and Horwitz, 2018). On the other hand, drawing samples in excitation space and then fitting them to the given LED stimulus system will always ensure that as much of the available color space is tested (Fig. S6D-F). Colors outside the system’s gamut will be clipped, but if the system is chosen to efficiently span color space, clipping can be reduced significantly. Since photoreceptor sensitivities always have some overlap, the photoreceptor axes are not completely independent as is the case with the spatial dimensions of width and height. Thus, clipping would even occur in a perfect stimulus system. Our *drEye* Python package includes various ways to efficiently draw samples within the gamut of a stimulation system to avoid clipping.

## 5 Dealing with uncertain spectral sensitivities

So far, we have assumed that the spectral sensitivities are uniquely described. However, measured sensitivities can vary depending on the experimental methods used (Salcedo et al., 2003; Sharkey et al., 2020). Furthermore, eye pigments and the optics of photoreceptors can change the effective sensitivities of photoreceptors (Hart, 2001; Sharkey et al., 2020). This can lead to uncertainty in the measured sensitivity of photoreceptors within the experimental conditions of interest. For example, recent measurements of fly photoreceptors show a shift in the peak of the rh6-expressing photoreceptor and a general broadening in the photoreceptors, as compared to original measurements (Sharkey et al., 2020). These differences likely reflect differences in sample preparation and measurement technique. Instead of having to (re-)measure the sensitivities within the experimental context of interest, previous measurements can be taken into account to build a prior distribution of photoreceptor sensitivities.

In order to take into account the distribution of photoreceptor properties, we first construct a normalized variance matrix **Σ**:

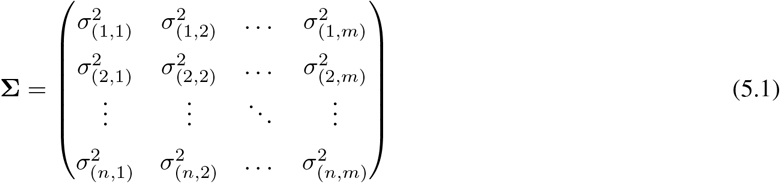

Here, 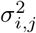 is the estimated variance of the relative light-induced capture of photoreceptor *i* given the light source *j* at an intensity of one unit photon flux. We can estimate each variance by drawing samples from the distribution of photoreceptor properties, then determining the light-induced capture for each sample given each light source at one unit photon flux, and finally calculating the empirical variance across samples. Given a particular set of light source intensities x, we can approximate the total variance of the calculated excitations *ϑ*^2^ by propagating **Σ** using Taylor expansions:

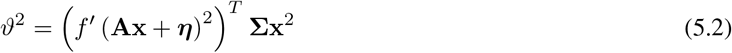

A large value of *ϑ*^2^ indicates that the chosen intensities x result in calculated excitations that vary considerably between different samples from the prior distribution of photoreceptor properties. We are less certain that the calculated excitations match the desired target values. Conversely, a smaller value of *ϑ*^2^ indicates that we are more certain that the calculated values match the desired target excitations. Thus, we wish to minimize *ϑ*^2^ while also matching the target excitations. To do this we apply a two-step procedure. The first step uses the mean spectral sensitivities and fits the excitation values as described in equation 3.7. We use the fitted x from the first step as our initial guess for the second step. In the second step, we minimize *ϑ*^2^, while constraining our solution for x to not deviate significantly from our fit in the first step:

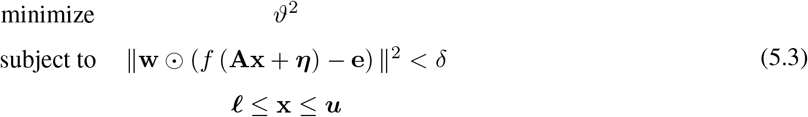

*δ* is the value of the objective function after optimizing x in the first step plus some added small value. *δ* may even be zero in an underdetermined system as multiple solutions can exist that give an optimal fit but have different values for the overall uncertainty *ϑ*^2^.

As an example, we consider two photoreceptors with spectral sensitivities that follow a Gaussian distribution (Fig. 4A) and an underdetermined stimulus system consisting of a UV, green and orange LED (Fig. 2G). The widths of the sensitivities vary between 30-70nm and 60-100nm for each photoreceptor respectively. The peaks of the sensitivities vary between 420-460nm and 500-540nm for each photoreceptor respectively. Changing the width and/or peak of the spectral sensitivity of either photoreceptor will affect the calculated capture for each LED differently. As an example of fitting using equation 5.3, we take a look at four different relative capture values (Fig. 4B). All four values are within the gamut of the stimulation system for the expected spectral sensitivities. As this is an underdetermined system, we can find multiple LED intensities x that fit the desired capture values for the expected sensitivities (Fig. 4C). Using only the standard fitting procedure from section 3.2, we find a set of intensities that has the smallest L2-norm (Xs in Fig. 4C). If we subsequently optimize to minimize the variance *ϑ*^2^, the fitted intensities x can differ significantly from the first fitting procedure (open squares in Fig. 4C). However, the overall goodness-of-fit for the expected target excitations is not affected as the solution simply moved within the space of possible optimal solutions (lines in Fig. 4C). To get a better idea of how fits differ between the original approach and this variance minimization approach, we drew many random capture values that are inside and outside the system’s gamut (Fig. 4D). After fitting, we calculated the average *R*^2^ scores of the two approaches for all samples from the prior distribution of spectral sensitivities. We find that on average the variance minimization approach improves the *R*^2^ score when considering a distribution of possible spectral sensitivities (Fig. 4E).

**Figure 4.**
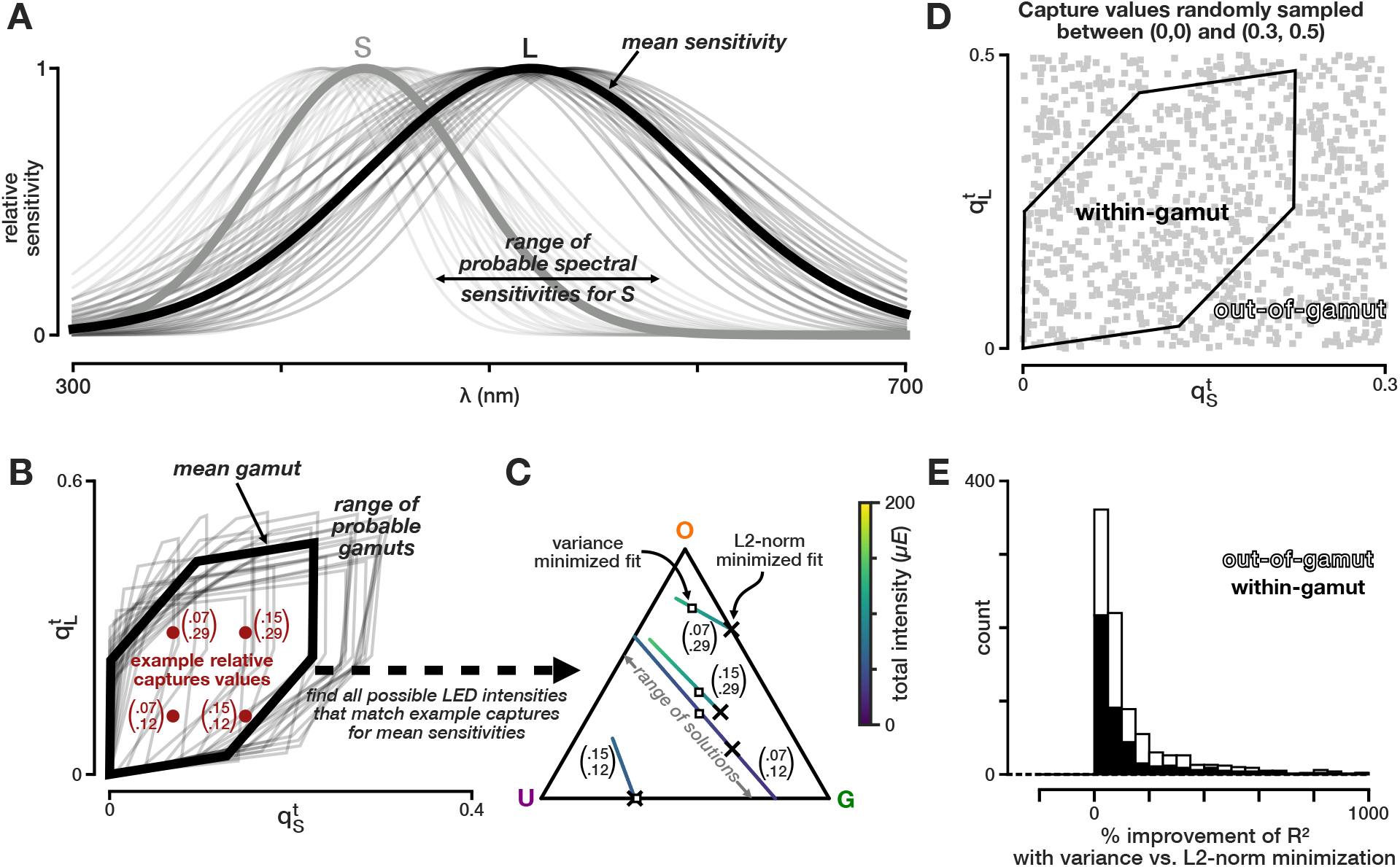
Minimizing the variance in excitation values due to uncertainty improves the average fit. **(A)** Variance in the spectral sensitivity for the short and long photoreceptor from the example in figure 2B by varying the mean between 420-460nm and 500-540nm for each photoreceptor in steps of 10nm and varying the standard deviation between 30-70nm and 60-100nm in steps of 10nm for each photoreceptor, respectively. **(B)** Relative capture space of the photoreceptors in A adapted to a flat background spectrum. Gamut of the LED set in figure 2G (thick line) and the resulting variance of the gamut due to the variance in the spectral sensitivities (thin lines). Xs correspond to example capture values that are within the gamut given the expected sensitivities in A (thick lines). **(C)** Possible LED proportions that result in the same calculated capture for the four examples in B using the expected sensitivities and the stimulation system from figure 2G. Each colored line corresponds to the set of proportions that result in the same capture. The color indicates the overall intensities of the set of LEDs. Xs indicate the fitted LED intensities using the fitting procedure defined by equation 3.7. The open squares indicate the fitted intensities after minimizing the variance according to equation 5.3 given the uncertainty in the spectral sensitivities as shown in A. **(D)** Randomly drawn captures that are in- and out-of-gamut (gray squares). **(E)** Average improvement in the *R*^2^ score for all possible samples of the spectral sensitivities in A when fitting the points in D with the additional variance optimization step. The black bars correspond to within-gamut samples and open bars correspond to out-of-gamut samples.

The variance minimization approach works well when the goal is to fit particular excitation values (e.g. to span the excitation color space as in section 4.2). However, if the goal is instead to fit particular spectral distributions, the method can lack accuracy because the corresponding target excitation values are not unique due to uncertainty of photoreceptor sensitivities. To deal with this problem, we can increase the number of photoreceptors we use to fit artificially by adding different samples of sensitivities to the estimation procedure and weighting them by their prior probability. We can still perform variance minimization as a second step, if the sampled sensitivities cover a range of possible excitation values.

A final approach to dealing with uncertainty of the spectral sensitivities is to update the prior distribution of the sensitivities with different behavioral and/or physiological data. Besides the more classical approaches to assessing the spectral sensitivity of photoreceptors and the dimensionality of color vision, sets of metameric stimuli can be designed to probe the responses of various neurons responding to visual inputs. For example, the lines in figure 4C correspond to a range of metameric stimuli that match the four example target captures in figure 4B. In the *drEye* package, we provide various ways to design metameric stimuli. For another example, we have measured the responses of photoreceptor axons in the fruit fly to stimuli that simulate the spectrum of one LED and then compared this response to the response of the neuron to the actual LED (Fig. S7). If the responses match, the sensitivities used are a good approximation within the wavelength range tested. However, care should still be taken as different sensitivities can still produce the same response in a (randomly) chosen neuron or behavior. Thus, many different types of neurons should be measured and stimuli tested for validation purposes.

## 6 Application to patterned stimuli

So far we have not explicitly considered the spatial aspect of chromatic stimuli. Our method can be used to display not only full field stimuli but also patterned stimuli, simply by applying it pixel by pixel. However, specific considerations need to be taken into account when it comes to these types of stimuli, which depend both on the animal and the hardware.

Our method can directly be applied to LCD screens and any other screens that pack small LEDs onto single pixels (Powell et al., 2021), as long as the set of light sources are adequate for the animal in question (see section 3.3). However, if the gamut afforded by the available set of LEDs used in a particular display is small, it will require a change in the LED set *at every pixel,* a time consuming and expensive task. In such cases, projectors offer a more flexible and affordable solution, as these only require replacing a single set of light sources by either swapping one or more LEDs or filters or using fiber optics to couple an external light source (Bayer et al., 2015; Franke et al., 2019). However, the effective use of our method in the context of this type of hardware depends on two factors: the dimensionality of color vision and the flicker fusion rate of the animal of interest.

Indeed, the core principle of projector design relies on “temporally” mixing light sources in different ratios in repeating patterns of subframes, and doing so at a higher frequency than the flicker fusion rate of the viewer (Fig. 5A). In most modern video projectors, this mixing occurs independently at each pixel, owing to an array of mirrors that are synced with each subframe and thus allow for a patterned image to be formed. This method at its core is equivalent to the algorithm we presented above, mixing in time instead of space.

**Figure 5.**
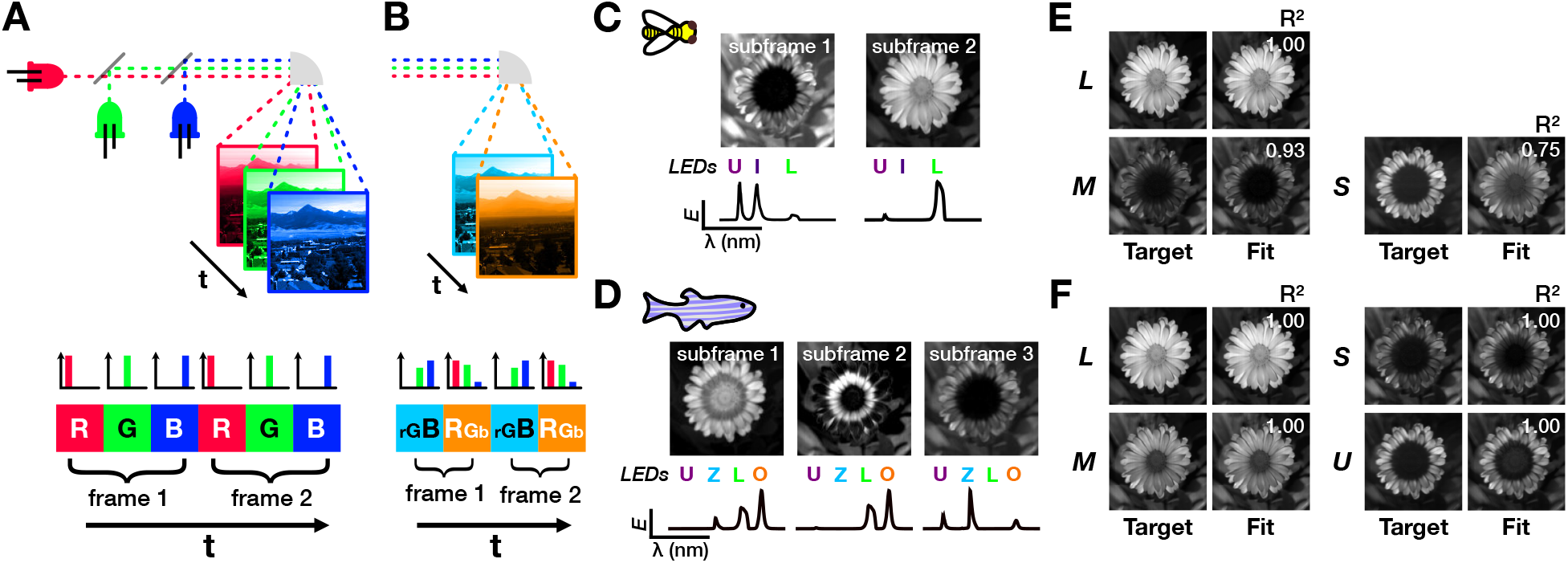
Reconstructing hyperspectral images with fewer subframes and number of photoreceptor types in the honeybee and zebrafish. **(A)** Schematic of the subframe structure in traditional RGB projectors **(B)** Schematic of a subframe structure with fewer subframes than LEDs. **(C-D)** Reconstruction of a hyperspectral flower (Resonon Inc.) in the honeybee and zebrafish with two or three subframes and three or four LEDs, respectively. The top images are the 8-bit mask for each subframe and the bottom are the normalized LED intensities used for each subframe. **(E-F)** Comparison of target photoreceptor captures and fitted captures for each photoreceptor type for the honey bee and zebrafish, respectively. The *R*^2^ value for each photoreceptor type is indicated in the image of fitted values.

Importantly, in such systems, *each subframe is dedicated to one light source.* Therefore this subframe structure typically limits the experimenter to use only up to a number of independent light sources equal to the number of subframes, to reconstruct a light spectrum at each pixel. If this number is equal or larger than the number of photoreceptor types of a given animal, and if the refresh rate of the hardware is higher than the flicker fusion rate of the animal, the algorithm detailed above can be applied to reconstruct patterned images. However, when both of these conditions are not met, the method is not suitable. If there are fewer subframes than photoreceptor types, the gamut of the system will be too small to properly reconstruct most images. If the flicker fusion rate of the animal is higher than the refresh rate of the hardware, the temporal mixing will not work, and the subframes will be seen as flickering.

For such cases, we have instead developed a different algorithm that can alleviate either problem, by allowing the use of a higher number of light sources than subframes and using the high spatial-spectral correlations existing in natural images, to optimally mix light sources in each subframe. We take advantage of modifications to some projector systems which allow for a more flexible use of their subframe structure. This flexibility in practice lifts the requirement of one dedicated light source per subframe, giving the user control over the spectral composition of each subframe (Bayer et al., 2015; Franke et al., 2019).

As pixel intensity and light source intensities can be manipulated independently, the aim of our algorithm is to find the best light source intensities **X** (sources x subframes) and pixel intensities **P** (subframes x pixels) for each independent subframe, so that they match the target photoreceptor excitations **E** (photoreceptors x pixels) of the whole image:

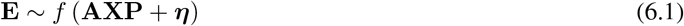

The light sources are constrained by the lower *ℓ* and upper **u** bound intensities they can reach, whereas the pixel intensities are parameterized, so that 1 allows all light from the light sources to go through and 0 does not allow any light to go through (i.e. luminosity). We fit **P** and **X** using an iterative approach similar to common EM-type algorithms, where we fix either **P** or **X** at each iteration while fitting the other using the same nonlinear least-squares approach as previously. To initialize both **P** and **X** we first decompose the relative capture matrix of the image **Q** (photoreceptors x pixels) using standard non-negative matrix factorization. This returns two non-negative matrices **P**_0_ (subframes x pixel) and **Q**_0_ (photoreceptors x subframes), whose dot product approximates **Q**. **P**_0_ is normalized so that its maximum is 1 and used as the initial matrix for **P**. For each column – i.e. each subframe – in **Q**0, we apply the nonlinear transformation to obtain excitation values and then fit light source intensities according to our objective function in equation 3.7. Using the initial values for **P** and **X** only a few iterations are needed (usually <10) to obtain a good fit for reconstructing the image.

As an example, we fit a hyperspectral image of a calendula flower (Resonon Inc.) given the bee and the zebrafish photoreceptor sensitivities and their corresponding optimal LED sets (Fig. 5). For both animals, we set the number of subframes to be smaller than the number of photoreceptor types (2 subframes for the trichromatic bee, and 3 subframes for the tetrachromatic zebrafish), allowing for a high projector frame rate and high image bit-depth to be set given the experimenter’s hardware. Despite having fewer subframes than photoreceptors, we are able to achieve good fits by mixing LEDs available in each subframe at different intensities, showing that we can effectively increase the refresh rate of a projector system for trichromatic animals, or use four LEDs for tetrachromatic animals without sacrificing our fits. It is important to note that, although this method works well in most cases, it may sometimes be impossible to achieve perfect fits for every photoreceptor, depending on given photoreceptor sensitivities and the spectral correlations of the hyperspectral image. An example of this is clear for the fitting of the S photoreceptor of the bee in our example image, only reaching a R^2^ value of 0.753. In such cases, hardware limitations may prompt the experimenter to use more subframes at the cost of the projector frame rate.

## 7 Conclusion

While studies in trichromatic primates have benefited from the wide adoption of consistent methods for designing chromatic stimuli, studies in other animals have suffered from a lack of uniform methodology. This has resulted in difficulties in comparing experimental results both within and between animals. More generally used chromatic stimuli – e.g. using monochromators or standard RGB displays – also do not take into account the color space of the animal under investigation and usually give an incomplete description of the properties of a color vision system. Furthermore, with the currently available techniques, it has been challenging to design more natural stimuli, especially natural images, and thus understand the role of spectral information in processing ecologically-relevant scenes.

Here, we present a method for designing chromatic stimuli, founded on color theory, that resolves these issues and can suit any animal where the spectral sensitivities of photoreceptors are known using a minimal visual stimulation system. Specifically, we provide a series of tools to reconstruct a wide range of chromatic stimuli such as targeted and random stimuli as well as hyperspectral images. We offer refinements to our methods to handle various nuances of color vision, such as uncertainty in spectral sensitivities or handling out-of-gamut color reconstruction. Even though our methods are hardware agnostic, we provide guidelines for assessing the suitability of a given stimulus system or selecting *de novo* light sources. Because our methods do not depend on the stimulation device itself, they can serve as a color management tool to control stimulus systems within and between laboratories and therefore improve reproducibility of experimental results.

In addition to the tools that we present here, our Python package *drEye* contains other tools that we have only mentioned briefly or omitted. These include efficient and even sampling of the available gamut, designing metameric pairs in underdetermined stimulation systems, and finding silent substitution pairs (e.g. section D). In addition, we have focused here on receptor spaces as a foundation for building stimuli, however if further transformations of receptor excitation, such as opponent processing, are known, these can also be included using our package, allowing the user to work in a space that might be “closer” to the animal’s perceptual space. Future updates will include new features such as the possibility of taking into account the varying spatial distribution of photoreceptor types across the eye of many animals (Wernet et al., 2015). With the aim of making adoption of our methods effortless, we provide our open-source *drEye* API and will make an accessible web application, which will be easy to use, regardless of coding proficiency.

## Code availability

The *drEye* Python package is available under **https://github.com/gucky92/dreye**, and a list of the essential dependencies are listed in Table S1. Tutorials for using the different methods mentioned in the paper and additional approaches that were ommitted can be found in the documentation for the package under **https://dreye.readthedocs.io/en/latest/**.

## Acknowledgments

We thank Larry F Abbott, Darcy Peterka, Elizabeth Hillman, and Sharon Su for comments on the manuscript. MPC was supported from NIH 5T32EY013933 and NIH R01EY029311. SLH acknowledges support from NSF GRF DGE-1644869 and NIH F31EY030319. SNM was supported by NIH R01EY029311. RB was supported by NIH R01EY029311, the McKnight Foundation, the Grossman Charitable Trust, the Pew Charitable Trusts, and the Kavli Foundation.

## Author contributions

MPC and RB conceived and designed the project. MPC developed the mathematical algorithms, designed, wrote, tested, and documented the software and performed analysis of the examples. MPC and SNM implemented and tested the algorithm for reconstructing patterned stimuli. SNM processed and analyzed the hyperspectral images. MPC, RB, SLH, and SNM wrote and revised the manuscript. SLH acquired the imaging data and performed fly husbandry. MPC processed and analyzed the imaging data.

## Supplementary Material

### A Projecting capture values into the chromaticity diagram

To project (relative) capture values of any set of photoreceptors onto the *n* — 1 simplex, we first need to normalize the *n* capture values by the L1-norm to obtain the porportional captures **p**:

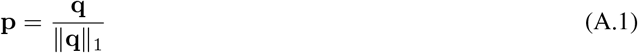

Next, we define a linear transformation **T** (*n* – 1 x *n*) that project **p** onto the *n* – 1 simplex:

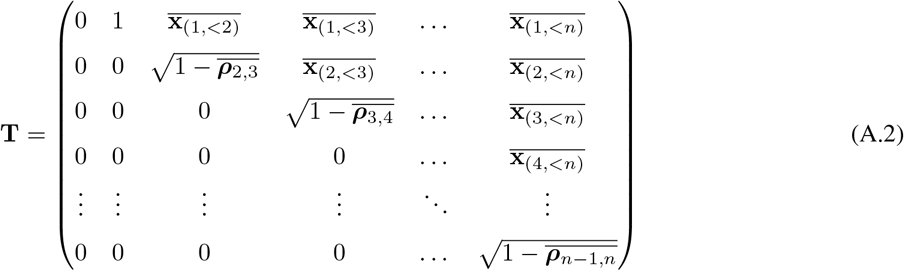

where 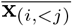 is the average for all values in row *i* before the *j*th column, and 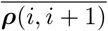 is the average of 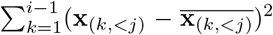 given row i and column *j = i* + 1. For example, 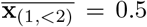 and 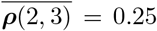. These formulations ensure the equality of all the distances between all the vertices in the *n* – 1 simplex. The point of capture values in the chromaticity diagram is then defined by **Tp**. We can also center the simplex by subtracting the point of equal capture from our capture values **Tp** — **T1**/*n*.

### B Calculating the percent of colors represented by a stimulation system

Our gamut metric – the percent of colors represented by a stimulation system – is the ratio of the mean width of the stimulation system in the chromatic space of an animal over the mean width of an ideal system in this space.

To calculate this metric we separately consider a “perfect” stimulation system, where the intensity of each unit wavelength along the (animal) visible spectrum can be varied independently, and the “real” stimulation system, composed of a combination of light sources. The intensity-independent gamut of the “perfect” stimulation system is the convex hull of the single wavelength manifold in the n — 1-dimensional chromaticity diagram (Fig. S2C, multi color line). To measure the size of this body (the convex hull), we calculate its mean width:

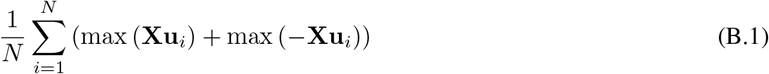

where **X** are the mean-centered vertices that describe the convex hull in the chromaticity diagram (points x chromaticity dimension) and **u**_*i*_ is one of *N* L2-normalized random vectors drawn from a standard normal distribution. We drew a total of 10000 random vectors (*N* = 10000). We also project a set of possible captures of the stimulation system in question (the “real” one) onto the chromaticity diagram S2C, gray shape). The set of captures we include are the calculated captures from all possible combinations of turning the LEDs off or maximally on. This will include all vertices of the convex body of the “real” stimulation system. As with the “perfect” system, we calculate the mean width of “real” stimulation system. The ratio of the mean widths corresponds to the fraction of colors that can be represented by a given stimulation system relative to a “perfect system” for a given animal.

The sum of capture values – the overall capture – approximates the overall intensity of a stimulus. Before projecting the set of possible captures of a stimulation system onto the chromaticity diagram, we can linearly interpolate between points to obtain points at specific overall capture and thus obtain the gamut metric for different intensity regimes. Consequently, the percentage of colors that can be represented by a given stimulation system drops significantly for high intensity regimes (Fig. S2D-E). Unlike in the intensity-independent case, having more LEDs than the number of photoreceptors can significantly improve our gamut metric. Therefore, we can determine if and when adding additional LEDs would enable reconstruction of more colors at higher intensities.

### C Gamut corrections prior to fitting

More details on gamut corrections methods available in our package, can be found on the github page for *drEye*. Here is just a short summary of two methods to correct a set of capture values obtained from an image to prevent burning of the image.

#### C.1 Scaling of the overall capture values to fit within the range of intensities of the stimulation system

If the stimulation system does not reach the intensities required to accurately reconstruct a set of stimuli (i.e. excitation values), we can scale the target relative capture values **q** linearly for all stimuli. To do this we first multiply the normalized capture matrix **A** element-wise by the maximum possible intensity *u* of each light source:

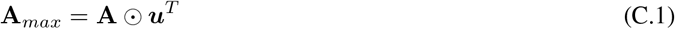

Then we calculate the minimum of the maximum in each row of **A**_*max*_:

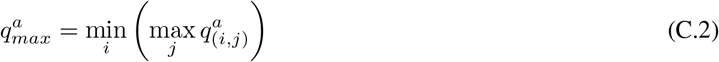

where 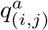 is the element in **A**_*max*_ corresponding to photoreceptor *i* and light source *j*.

Similarly, we calculate the maximum relative capture across the whole set of target stimuli:

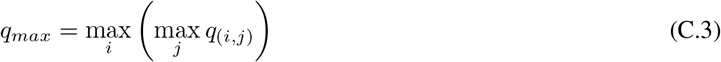

where *q*_(*i,j*)_ is the relative capture of photoreceptor *i* and stimulus *j*.

Finally, we scale all target capture values, so that they span the intensity range of the stimulation system that is able to reproduce the most colors:

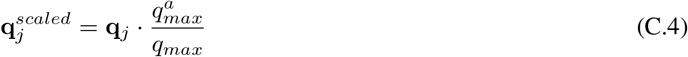

Using the rescaled relative captures, we can then apply the baseline term *η* and the nonlinear transformation *f* to obtain the rescaled target excitation values.

#### C.2 Scaling of capture values to fit within the gamut

Our fitting algorithm will automatically clip values that are outside of the system’s gamut. However, this simple clipping can burn a set of target values as the relative distances in receptor-based capture space are not preserved. After applying the transformation from the previous section, we can further rescale capture values, so that they are within the gamut of the stimulation system. That is all capture values are inside the body defined by the light sources of the stimulation system in the chromaticity diagram (e.g. gray shapes in Fig. S2C-G). To do this we first project all target captures onto the chromaticity plane (see section A) to obtain a matrix **P** (stimuli/pixels x photoreceptors). For humans, rescaling and clipping usually occurs to preserve hue while caring less for saturation of a color. This effect can be quasi-replicated in the chromaticity diagram for any animal by aiming to preserve the angles relative to the adapted capture – the center in the chromaticity diagram. To adjust the saturation of a stimulus, we need to adjust the distance of all capture values to the center of the chromaticity diagram until all points are within the gamut of the stimulation system. First, we find by how much each set of capture values has to be rescaled in order to fit within gamut. Then, we take the minimum scaling value and scale all capture vectors by that amount in the chromaticity diagram. This ensures that all points are within the gamut of the system. Finally, we project the capture values back into the receptor-based capture space and multiply them by their overall capture. Using the rescaled captures, we can then apply the baseline term ***η*** and the nonlinear transformation f to obtain the rescaled target excitation values.

In our *drEye* package, we also implemented an algorithm that performs the two described steps simultaneously using a convex optimization approach that finds the optimal trade-off between scaling the intensities and scaling the angles in the chromaticity diagram.

### D Silent substitution

The goal of silent substitution is to excite single (or a select set of) photoreceptor types while keeping all others silent. Silent substitution takes the spectral sensitivities of the chosen model organism into account in order to alter the stimulus in a way that only changes the excitation of a single or a subset of photoreceptors. More specifically, if the chosen stimulus presentation which activates the photoreceptor of interest also affects the excitation of other the photoreceptors, adjustments are to be made to negate those effects (Estévez and Spekreijse, 1982). While the silent substitution method is by no means new (Estévez and Spekreijse, 1982), here we offer a unified method that can be flexibly used regardless of stimulation system or model organism and does not require the experimenter to perform any calculations by hand.

The objective function for silent substitution is the following:

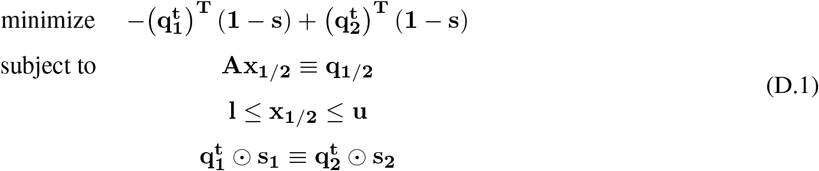

where *s* is an indicator vector with one indicating which photoreceptors to keep silent and 0 indicating the photoreceptors that are supposed to be differentially excited. The 1 and 2 subscript indicates the two stimuli/captures that aims to silence all photoreceptors but maximize the contrast for the photoreceptors that are not supposed to be silent. This objective function aims to find the maximum possible contrast between two stimuli for the non-silent photoreceptors given the LED system and spectral sensitivities of each photoreceptor.

For example, if an experimenter was attempting to isolate the blue-sensitive pR8 photoreceptor in the fruit fly, it would clearly require presentation of blue light to do so. However, this would also activate the blue-green-sensitive yR8. Thus, our method would also adjust the intensities of longer-wavelength LEDs to keep the response of yR8 the same, while only modulating the activity of pR8. Ultimately, this canonical method has been developed in a myriad of different settings, but our method unifies them by allowing any experimenter to customize the stimulus to their LED system and model organism specifics. Based on that information, our method then automatically finds the best LED combination that maximizes the contrast of the non-silent photoreceptors.

**Figure S1.**
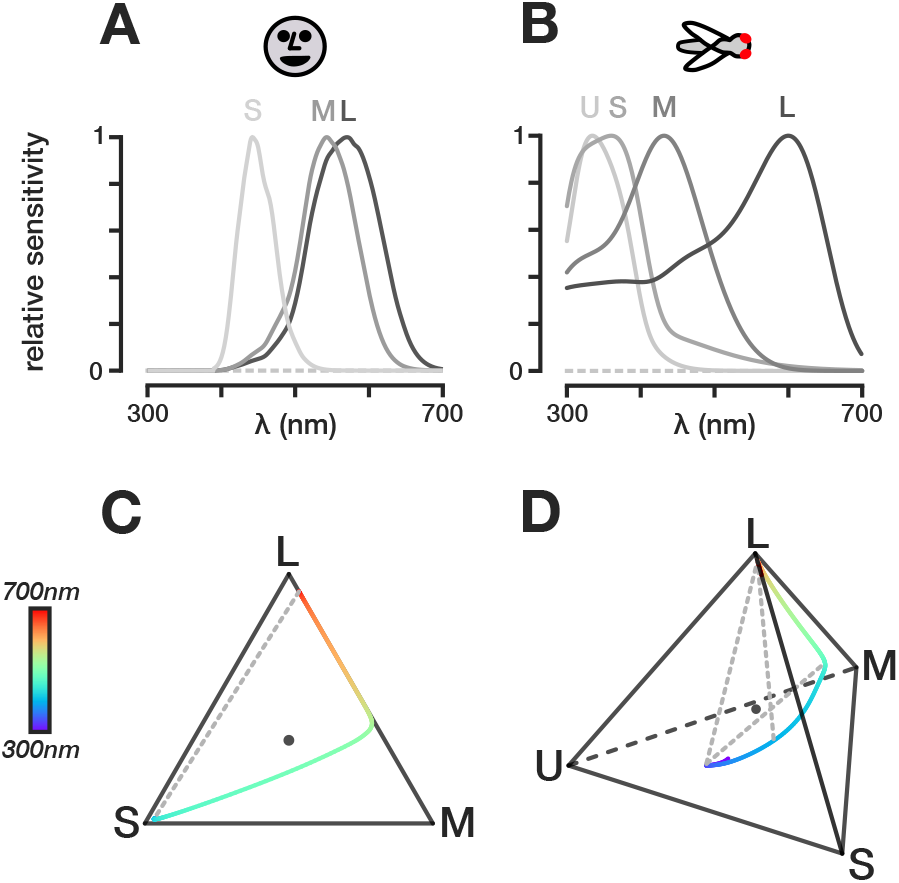
Sensitivities and chromatic diagrams of the human and fruit fly. **(A)** Human spectral sensitivities as measured by Stockman et al. (1993). **(B)** Spectral sensitivities of *Drosophila melanogaster* as measured by Sharkey et al. (2020). For simplicity, we ommitted the photoreceptor type expressing the rh1 opsin in the fruit fly as they are broadly sensitive to all wavelengths. **(C-D)** LMS chromaticity diagram for humans and fruit flies based on their spectral sensitivities, respectively.

**Figure S2.**
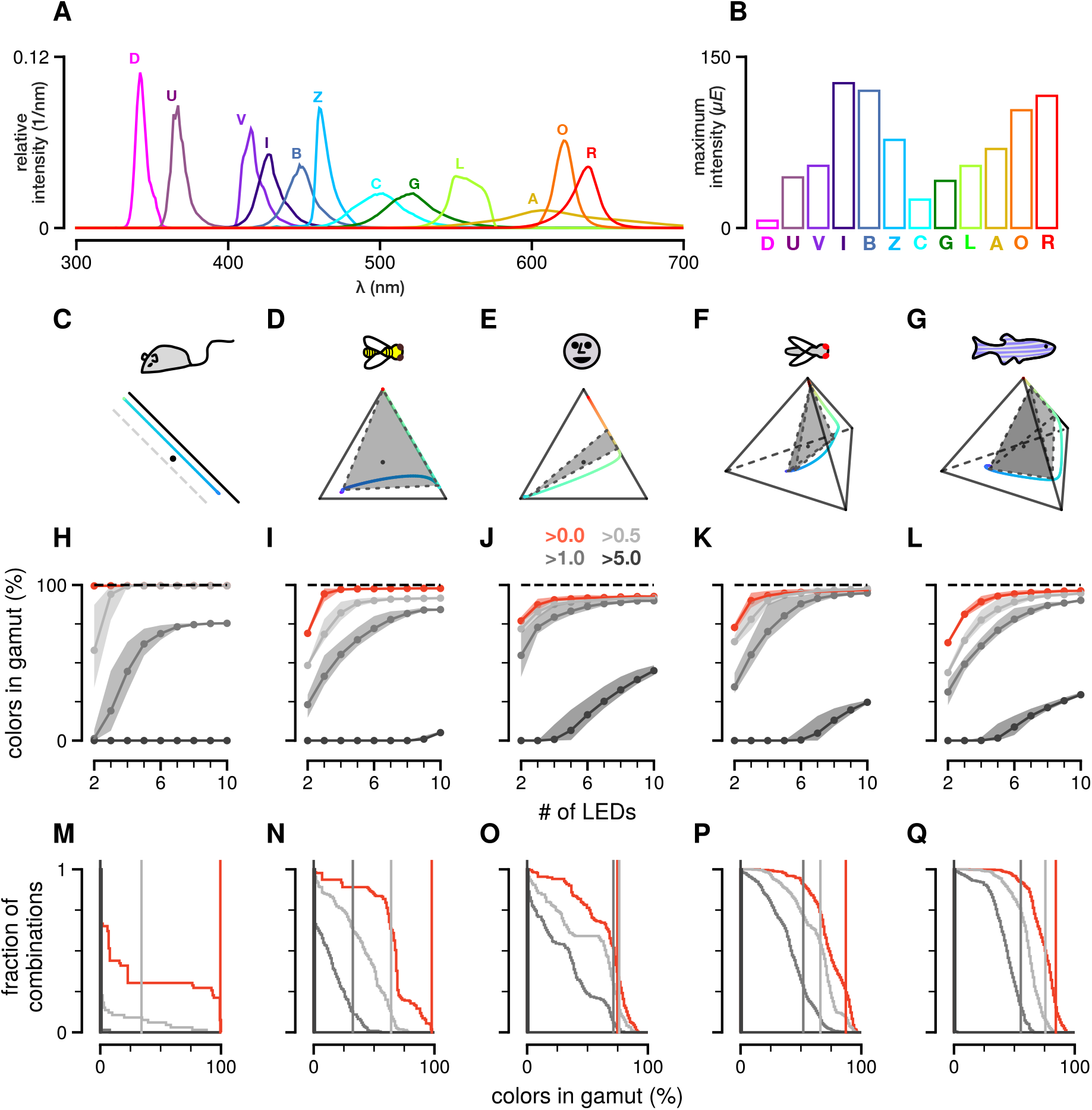
Color in the gamut across LED sets and animals. **(A)** Normalized photon flux across the wavelength spectrum for 11 LEDs currently available for purchase from ThorLabs. In order, the LED labels stand for: deep UV (D), UV (U), violet (V), indigo (I), blue (B), azure (Z), cyan (C), green (G), lime(L), amber (A), orange (O), and red (R). **(B)** Maximum intensity in photon flux *(μE* = *μmol/s/m*^2^) for LEDs in **(A)** calculated from the irradiance measurements provided by ThorLabs. **(C-G)** Chromaticity diagram of the mouse, honeybee, human, fruit fly, and zebrafish. The color line is the single wavelength line, which corresponds to the perfect system. The shaded area is the gamut of the LED stimulus system chosen according to the peak of the spectral sensitivities. **(H-L)** Percentage of colors in gamut for the best LED sets (top 10%) across different number of LED combinations for the mouse, honeybee, human, fruit fly, and zebrafish. Each line is the mean of the top 10% LED sets. The shaded area for each line spans the range of values for the top 10% LED sets. The red line is the percentage of colors in the gamut ignoring the intensity range of the LEDs. The gray lines are the percentage of colors in the gamut for different values of the overall capture. **(M-Q)** The cumulative distribution of the percentage of color in the gamut for all *n*-size stimulus systems. The vertical lines indicate the percentage of colors in the gamut for the LED set that most closely match the peak of the sensitivities.

**Figure S3.**
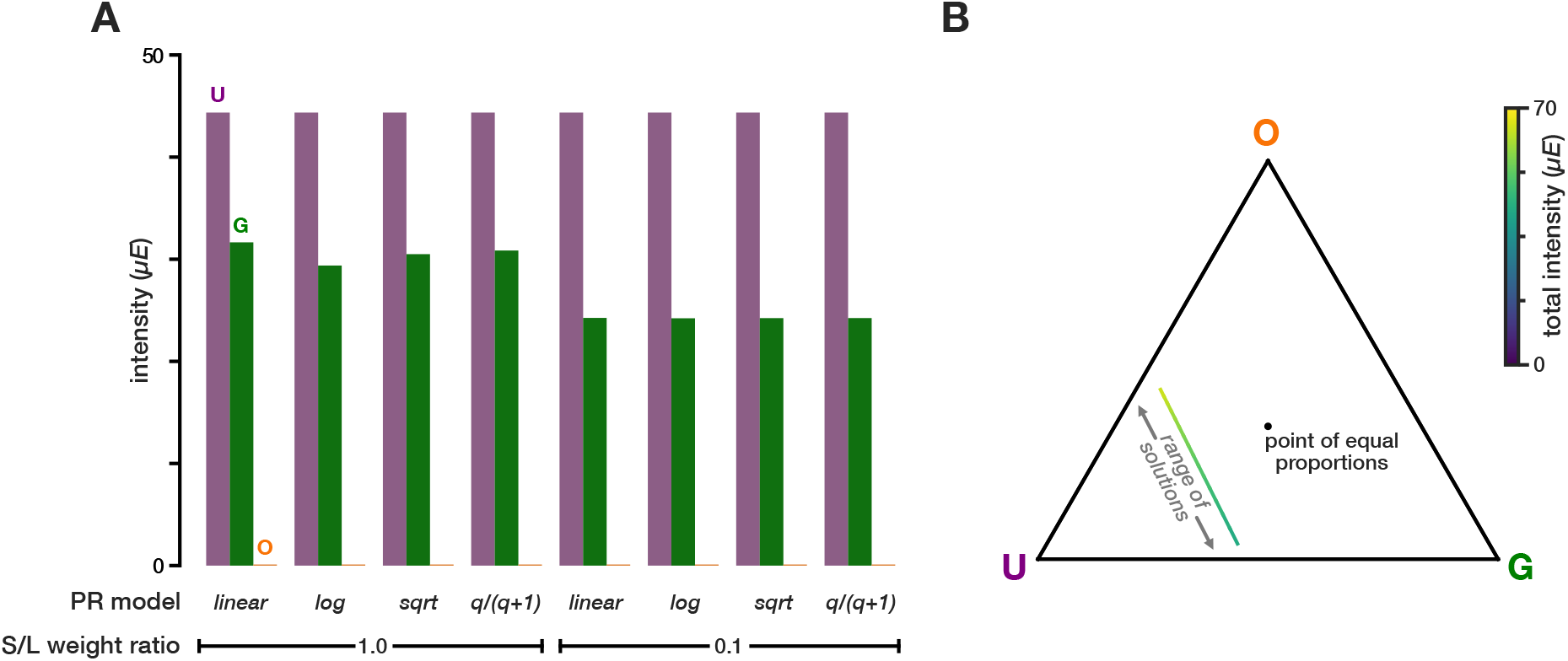
Fitting out-of-gamut spectra using different non-linearities and photoreceptor weights and fitting within-gamut intensities of underdetermined systems. **(A)** Fitting of the red-colored spectrum in figure 2A given the sensitivities in 2B and the stimulation system defined by the spectra in 2G using different non-linearities and photoreceptor weight ratios. In this case, using different non-linearities and weights mainly affects the intensity of the green LED. **(B)** Fitting the blue-colored spectrum in figure 2A given the sensitivities of 2B and the stimulation system in 2G yields many possible LED intensity combinations that give the same set of capture values (colored line). The color of the line indicates the overall intensity of the given intensity combination.

**Figure S4.**
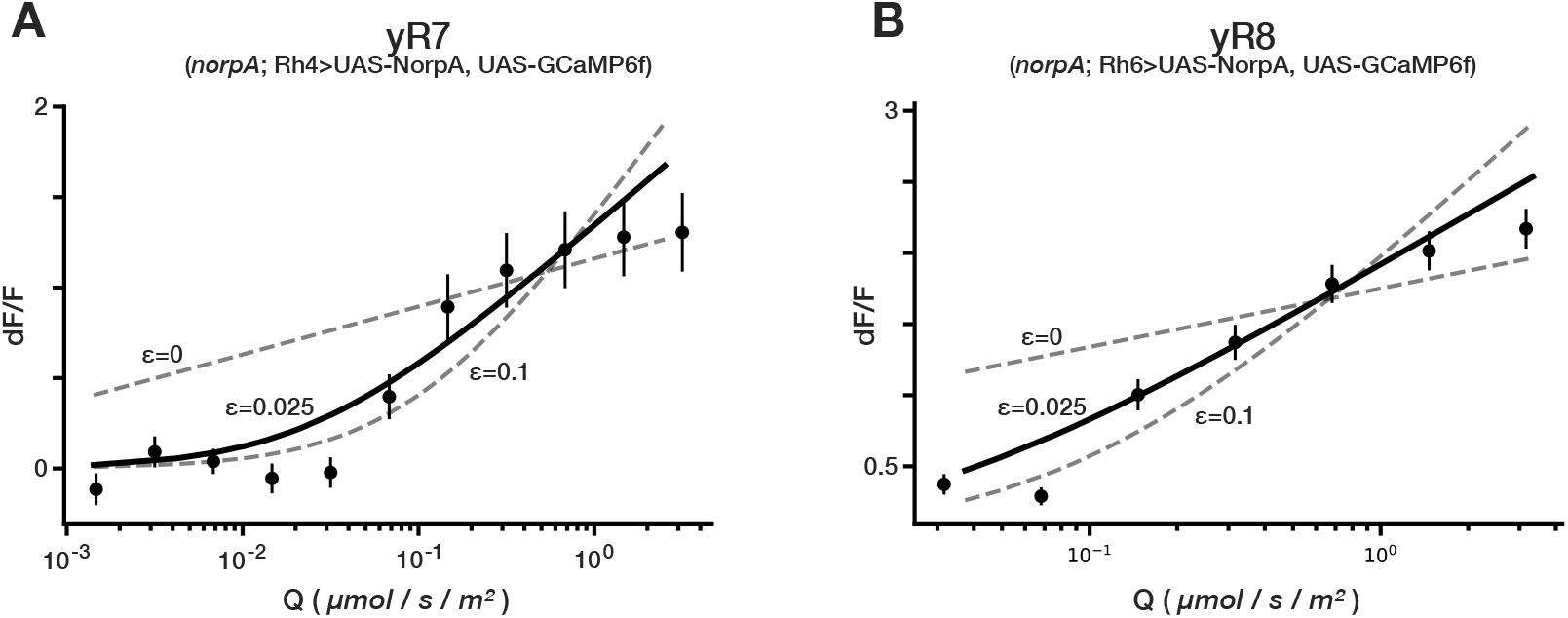
Responses of putative isolated photoreceptors to various combinations of lights fit a log photoreceptor model with a small baseline value *ϵ*. Two-photon calcium imaging was done as described in and using the same stimulation system as in Heath et al. (2020). The genetic fly lines are also the same *NorpA* lines used in Heath et al. (2020). The flies were adapted to the dark and different LED combinations were shown at an interval of 2 seconds with a duration of 0.5 seconds. The circles in panel A and B show the average response to all LED combinations that correspond to a small range of absolute capture values for the short-wavelength sensitive yR7 and long-wavelength sensitive yR8 photoreceptors, respectively. To calculate the capture values we used the spectral sensitivities as measured by Sharkey et al. (2020) (Fig. S1B). The error bars indicate the 95% confidence interval calculated as described in Heath et al. (2020). The solid line is the best fit for the baseline *ϵ* value given a log transformation of the relative capture values. In the dark, the relative capture values is calculated as follows: *q* = (*Q* + *ϵ*)/*ϵ*. To prevent zero division, *ϵ* = 0 is clipped to 10^-8^. The dashed lines are two different values for *ϵ* as indicated. yR7: 4 flies and 51 neurons; yR8: 6 flies and 120 neurons.

**Figure S5.**
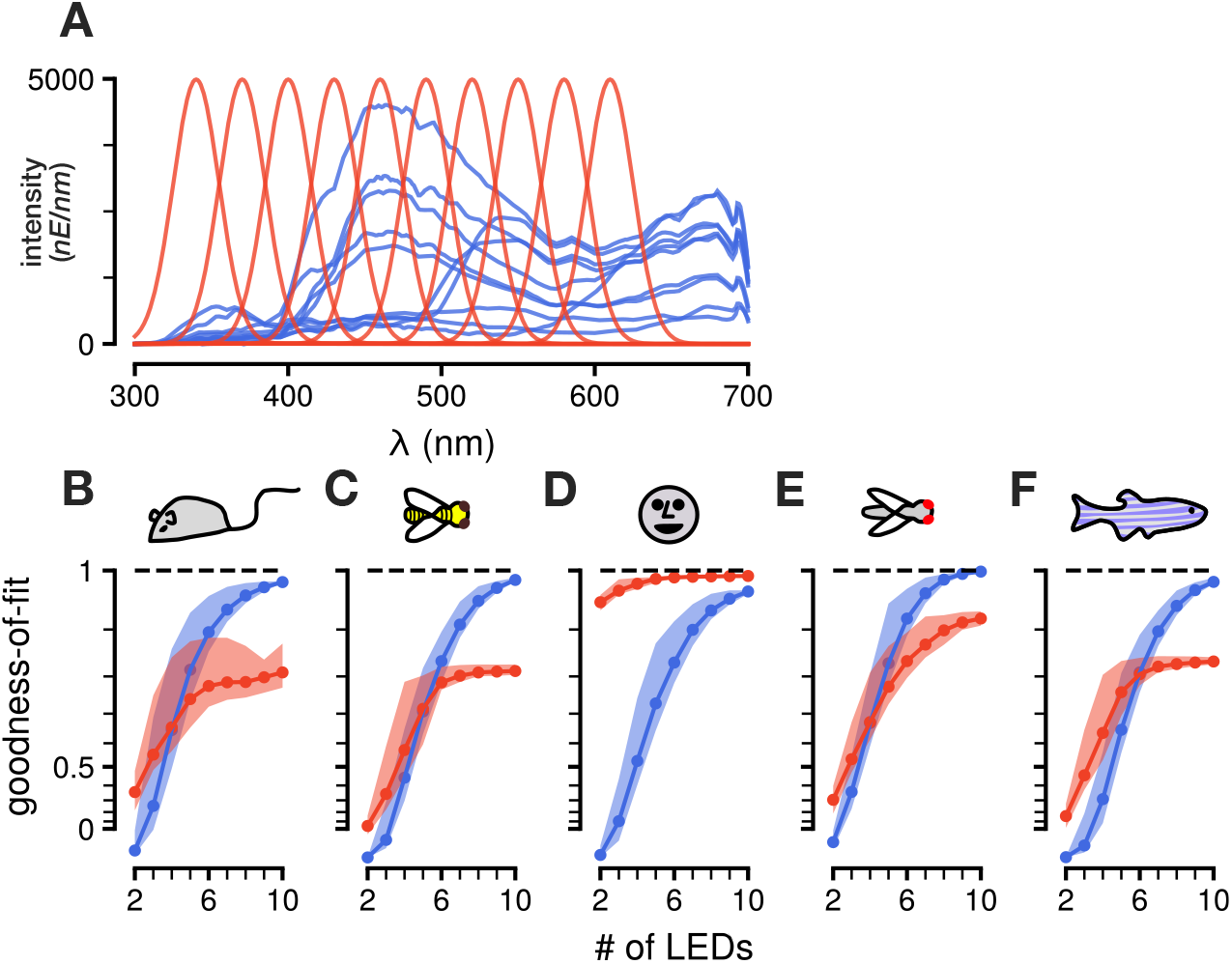
Fitting targeted stimuli to different model organisms at high intensities. **(A)** Example target spectra to be reconstructed at higher intensity than in figure 3 (same set as in figure 3A, but at 2000x the intensity): a set of natural spectral distributions (blue) and a set of Gaussian spectral distributions (red). **(B-F)** Goodness of fit (R^2^) values for the best LED sets (top 10%) across different number of LED combinations for the mouse, honey bee, human, fruit fly, and zebrafish, respectively. The filled areas correspond to the range of *R*^2^ values achieved for the top 10% LED sets.

**Figure S6.**
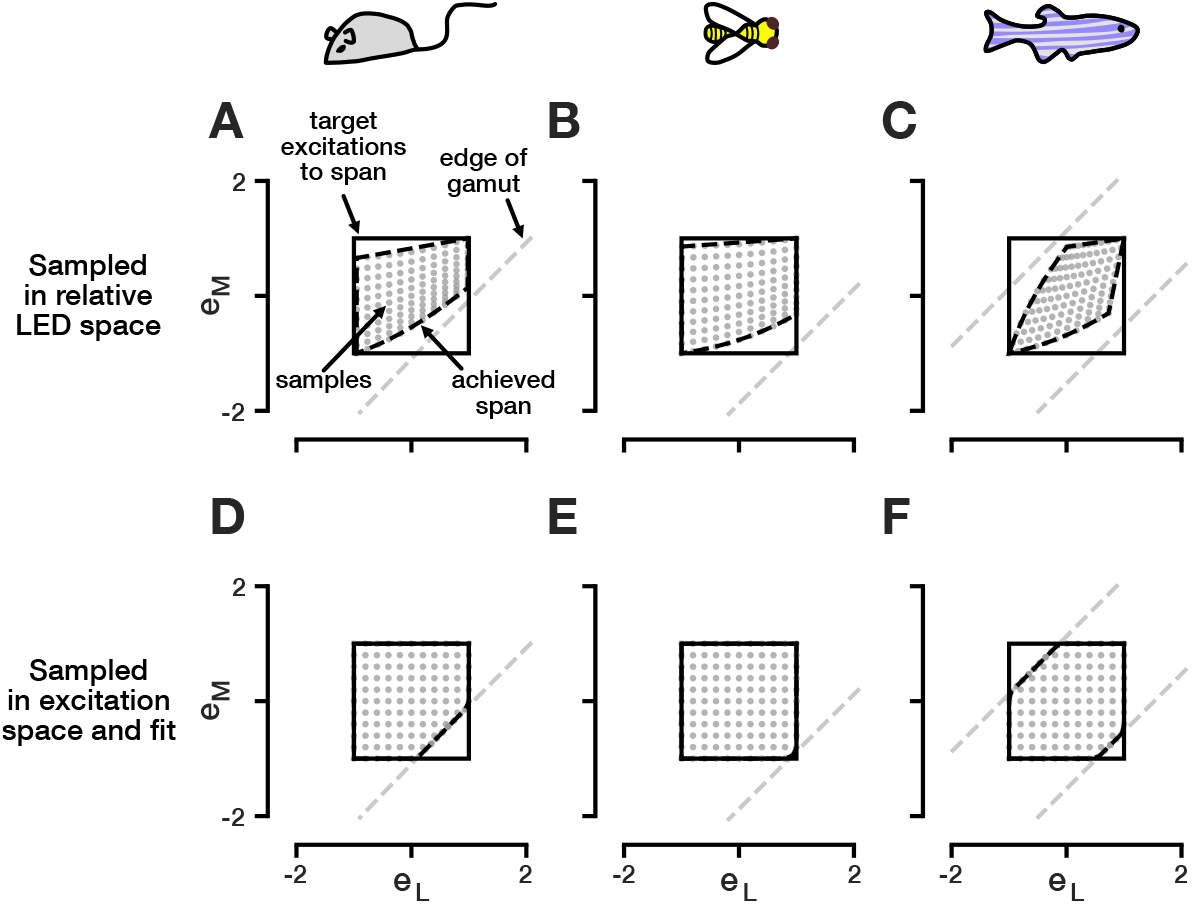
Fitting random stimuli to different model organisms. **(A-C)** Samples drawn in relative log LED space using the violet and lime LED (Fig. S2A) mapped onto excitation space of the mouse, honey bee, and zebrafish, respectively. The set of samples are more correlated along the achromatic direction (sum of excitations) than the chromatic direction (difference of excitations). **(D-F)** Samples drawn in excitation space and fitted using our optimization approach in equation 3.7. The set of samples span the chromatic and achromatic dimensions more equally within the limits of gamut of the stimulation system.

**Figure S7.**
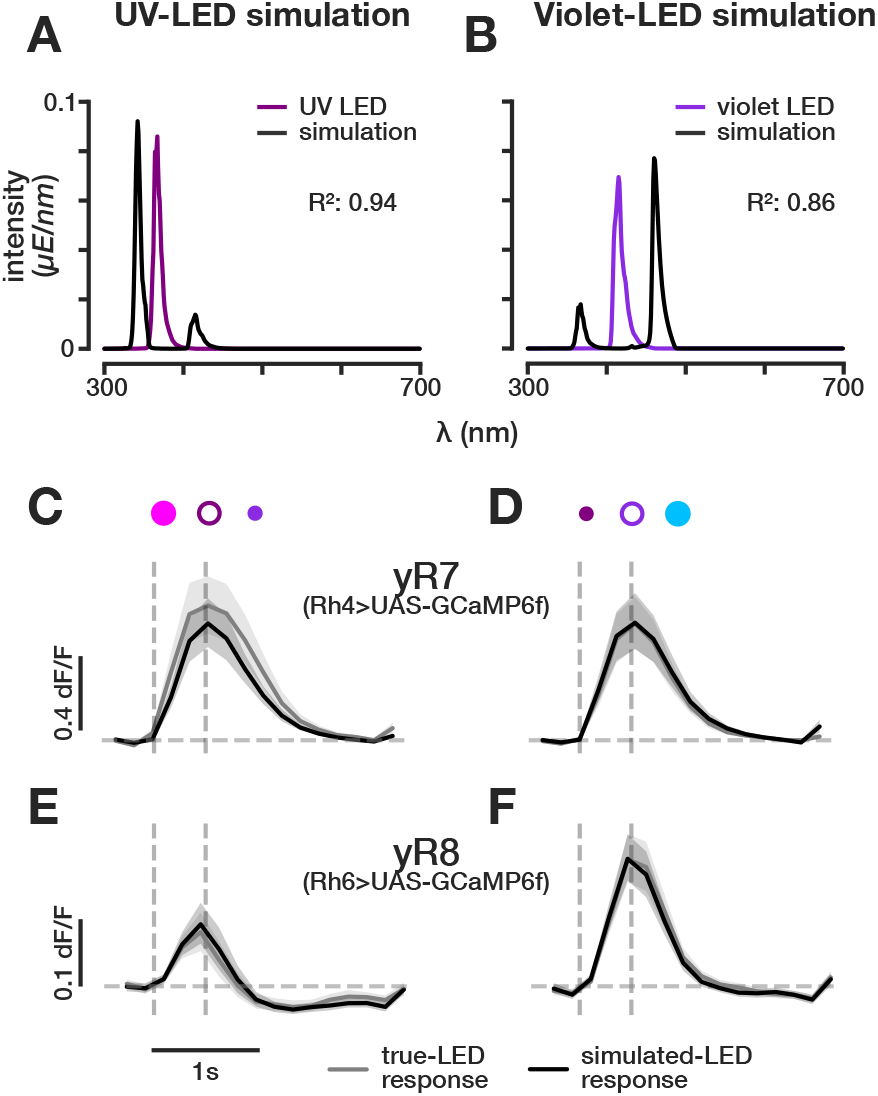
Performing LED simulations to verify previously measured sensitivities within the UV- and violet-wavelength range. Two-photon calcium imaging was done as described in and using the same stimulation system as in Heath et al. (2020). The genetic fly lines are also the same lines used in Heath et al. (2020). **(A-B)** Simulating the UV LED and violet LED using the surrounding LEDs: dUV and violet for UV LED simulation and the UV and azure LED for the violet LED simulation. We used the fruit fly sensitivities as measured by Sharkey et al. (2020) to fit the LED intensities for the simulations. **(C-D)** Responses of yR7 photoreceptor axons to the actual LEDs and their simulations. **(E-F)** Responses of yR8 photoreceptor axons to the actual LEDs and their simulations.

**Table S1.** Essential Package Dependencies of *drEye.*

| **Package** | **Version** |
| --- | --- |
| Python | $\geq 3.6$ |
| SciPy | $\geq 1.6$ |
| cvxpy | $\geq 1.1.10$ |
| quadprog | $\geq 0.1.9$ |
| jax | $\geq 0.2$ |
| scikit-learn | $\geq 1.0$ |

